# Diverse homeostatic and immunomodulatory roles of immune cells in the developing mouse lung revealed at single cell resolution

**DOI:** 10.1101/2020.02.10.942359

**Authors:** Racquel Domingo-Gonzalez, Fabio Zanini, Xibing Che, Min Liu, Robert C. Jones, Michael A. Swift, Stephen R. Quake, David N. Cornfield, Cristina M. Alvira

## Abstract

At birth, the lungs experience a sudden transition from a pathogen-free, hypoxic, fluid-filled environment to a pathogen-rich, rhythmically distended air-liquid interface. While many studies focus on adult tissue, the heterogeneity of immune cells in the perinatal lung remains unexplored. Here, we combine single cell transcriptomics with *in situ* hybridization to present an atlas of the murine lung immune compartment during a critical period of lung development. We show that the late embryonic lung is dominated by specialized proliferative macrophages with a surprising physical interaction with the developing vasculature. These macrophages disappear after birth and are replaced by a complex and dynamic mixture of macrophage subtypes, dendritic cells, granulocytes, and lymphocytes. Detailed characterization of macrophage diversity revealed a precise orchestration of five distinct subpopulations across postnatal development to fill context-specific functions in tissue remodeling, angiogenesis, and immunity. These data both broaden the putative roles for immune cells in the developing lung and provide a framework for understanding how external insults alter immune cell phenotype during a period of rapid lung growth and heightened vulnerability.

## Introduction

Prior to birth, the lung is maintained in a fluid-filled, immune-privileged, hypoxic environment. Upon birth, the tissue quickly transitions to an air-filled, immune-challenged, oxygen rich environment following the infant’s first breath^1^. At this point, the distal, gas-exchanging alveoli are suddenly subjected to the mechanical forces of spontaneous ventilation and exposed to diverse pathogens present in the external environment^2^. Adaptation to this rapid environmental shift is necessary for perinatal survival and is mediated by complex physiologic processes including reduced pulmonary arterial pressure, an exponential increase in pulmonary blood flow, establishment of air-liquid interface, and surfactant production^3^. The immune system is essential for lung homeostasis, wound-healing and response to pathogens^4^. Although the development of the murine immune system begins during early embryogenesis, little is known regarding how the dynamic physiologic changes at birth alter the lung immune cell landscape, and whether specific immune cell subpopulations influence lung growth and remodeling in addition to serving established immunomodulatory functions.

Immune cells play a central role in the development of many organs. Innate and adaptive immune cells regulate epithelial architecture during mammary gland development by promoting terminal end bud elongation and impairing ductal invasion^5^. Lymphocytes play a key role in oligodendrogenesis and synapse formation^6^, and macrophages inform kidney^7^, brain^8^, and retina^9^ organogenesis. In highly vascularized organs, macrophages localize to the tips of vascular sprouts to enhance vascular network complexity^9^, promote angiogenesis^10^, and regulate vascular patterning^11^. Although proximal lung branching occurs during early gestation, the development of distal airspaces capable of gas exchange begins only just before birth during the saccular stage of development. These saccules are subsequently divided into millions of alveoli after birth during alveolarization, the final stage of development characterized by rapid lung parenchymal and vascular growth^12^. Whether temporal regulation of specific immune populations informs lung immune function or the significant pulmonary parenchymal and vascular growth and remodeling occurring during early postnatal life remains unknown.

The prevailing notion is that the neonatal immune compartment is immature^13^. Limited immune competence, including attenuated innate immunity^14^, poor immuno-stimulatory function of antigen presenting cells^15^, and skewed adaptive immune responses may underlie the heightened susceptibility of infants to viral and bacterial infections^13^. Although the neonatal immune system can be induced to manifest adult-like responses under certain conditions^16^, this type-2-skewed immune environment likely facilitates immuno-surveillance and metabolic and tissue homeostasis^17^. Given that organ development occurs as the immune cell landscape is rapidly evolving, identifying the specific immune subpopulations present at discrete time points is vital to inform how immune cell diversity, localization, and cell-cell interactions may influence lung function and structure during early postnatal development.

In this report, we combined single cell transcriptomics (scRNA-Seq) with fluorescent multiplexed *in situ* hybridization (FISH), and flow cytometry to characterize changes in composition, localization, and function of immune cells in the murine lung from just before birth through the first three weeks of postnatal life. At birth, immune cell heterogeneity increased dramatically from an embryonic landscape dominated by immature, proliferative macrophages to a complex landscape comprised of multiple types of macrophages, dendritic cells, granulocytes, and lymphocytes. Dynamic changes in macrophage heterogeneity were particularly striking, both transcriptionally and spatially, including the presence of embryonic macrophages encircling developing vessels prior to birth. After birth, these embryonic macrophages disappear, and numerous unique macrophage populations emerge, each exhibiting unique gene signatures suggesting specific roles in immunosuppression, pathogen surveillance, angiogenesis, and tissue remodeling. Multiple populations of dendritic cells, basophils, mast cells and neutrophils are also present in the postnatal lung, expressing genes important for rapid pathogen response. In contrast, although lymphocytes increase in abundance across three weeks, they remain functionally immature and skewed toward type-2 immunity. Taken together, our data demonstrate a previously underappreciated plasticity of immune cells in the perinatal and neonatal lung suggesting unique and essential roles in regulating immune function and lung structure.

## Results

### Diversity of the lung immune landscape increases dramatically after birth

To comprehensively define the lung immune landscape at birth, we isolated whole lungs from C57BL/6 (B6) mice at four stages of perinatal development: the early saccular (E18.5), late saccular (P1) early alveolar (P7) and late alveolar stages (P21) (Figure 1A), and quantified gene expression by scRNA-Seq. Lung tissue was isolated, the pulmonary vasculature perfused to remove circulating immune cells, and the tissue digested using an in-house optimized protocol to ensure maximal cell viability as published protocols^18, 19^ induced high amount of cell death in the embryonic and early postnatal lung (Supplemental Figure 1). Live CD45+ cells were sorted by FACS, processed by Smart-seq2 and sequenced on Illumina NovaSeq 6000 (Figure 1A). Gene expression was computed as previously described^20^ and over 4,000 cells from 8 mice, 1 female and 1 male for each time point were analyzed, with an average of 1.03 million mapped read pairs and ∼3000 genes per cell (see Methods). To quantify whether the different mice contributed spurious variation to the data, a distribution level approach was chosen. For each cell type and time point, 100 pairs of cells from either the same mouse or different mice were chosen and the distance in tSNE space calculated. The cumulative distributions for those pairs were subsequently plotted to check whether pairs from different animals had a significantly longer distance than cells from the same mouse. We found no difference in the cumulative distributions (as also evident, on a qualitative level, by observation of the embeddings), indicating the absence of significant variation between the mice at each timepoint.

**Figure 1:**
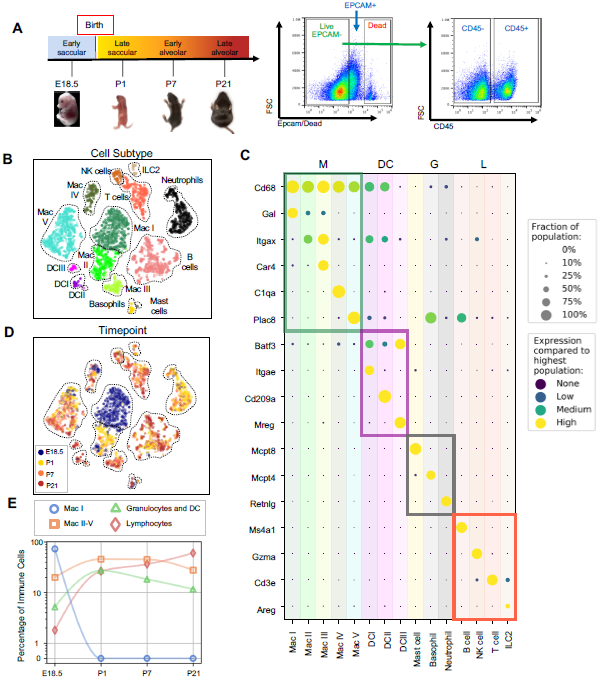
Diversity of the lung immune landscape increases dramatically after birth. (A) Overview of the experimental design including the four timepoints (E18.5, P1, P7, P21) corresponding to key stages in late lung development. Lungs were isolated, perfused, and digested and immune cells isolated by fluorescence activated cell sorting (FACS) for the dead-stain-, EPCAM-, CD45+ population. (B) t-Distributed Stochastic Neighbor Embedding (t-SNE) and unsupervised clustering of all immune cells identifies fifteen distinct populations. (C) Dot plot showing level of expression (purple to yellow), and fraction of the population expressing the particular gene (dot size) for distinguishing genes expressed by the Leiden clusters broadly separated into myeloid (M), dendritic cell (DC) granulocyte (G) and lymphocyte (L) populations. (D) t-SNE of immune cell clusters identifying developmental timepoint of cell origin with E18.5 (blue), P1 (yellow), P7 (orange) and P21 (red). (E) Quantification of the abundance of specific immune subpopulations in the lung at each developmental timepoint expressed on a log^10^ scale as percentage of total immune cells.

Fifteen cell clusters were identified via Leiden community detection^21^ and verified by t-distributed stochastic neighbor embedding (t-SNE)^22^ (Figure 1B). Myeloid cells separated into eleven clusters, including five distinct macrophage/monocyte subpopulations with shared expression of *Cd68*, and distinguished by expression of *Gal* (Mac I), *Itgax* (Mac II), *Car4* and *Itgax* (Mac III), *C1qa* (Mac IV), or *Plac8* (Mac V). Dendritic cells (DCs) separated into three clusters, all expressing some amount of *Batf3*, but distinguished by the expression of *Itgae* (DCI), *Cd209a* (DCII), or *Mreg* (DCIII). We also identified mast cells (expressing *Mcpt4*), basophils (*Mcpt8*), and neutrophils (*Retnlg*). Four lymphoid clusters were found, consisting of B cells (expressing *Ms4a1*), T cells (*Cd3e*), natural killer (NK) cells (*Gzma*), and group 2 innate lymphoid cells (ILC2) (*Areg*) (Figure 1C).

We next assessed cluster distribution across time (Figure 1D and 1E). Mac I cells dominate the late embryonic lung, with fewer macrophages scattered among clusters II, IV and V and an even smaller number of granulocytes and lymphocytes. After birth, immune cell heterogeneity increased explosively, concomitant with the disappearance of Mac I. Granulocyte abundance peaked just after birth and lymphocytes abundant increased progressively.

### Expression of *Dab2* and *Plac8* broadly separates macrophages and monocytes

Clusters Mac I-V exhibited the most striking heterogeneity, so we analyzed their transcriptomes and spatial distribution in detail. All five clusters shared high expression of *Cd68*, indicative of macrophages or monocytes (m/m) (Figure 2A). At E18.5, Mac I comprised ∼80% of m/m cells and Mac II, IV, and V each comprised 5-10% while Mac III was almost absent (Figure 2B and C). After birth, Mac I disappeared while Mac II abundance peaked to 35% of the total before decreasing again and disappearing by P21. Mac III and Mac V abundance increased steadily with Mac V being most abundant at all postnatal timepoints. The Mac IV population was relatively stable over time at ∼10% of the total.

**Figure 2:**
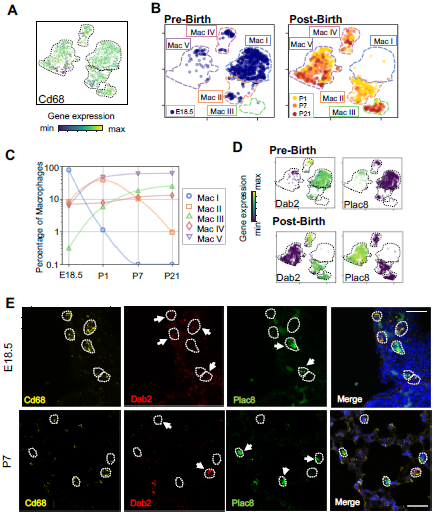
Macrophage populations present before and after birth broadly separate into two populations based on expression of *Dab2* and *Plac8*. (A) t-SNE plot depicting *Cd68* expression in the five macrophage populations. (B) Separate embeddings for prenatal versus postnatal macrophages, identifying developmental timepoint of cell origin with E18.5 (blue), P1 (yellow), P7 (orange) and P21 (red). (C) Quantification of the abundance of each macrophage subpopulations at each developmental timepoint expressed on a log^10^ scale as percentage of total macrophages. (D) t-SNE plots depicting expression of *Dab2* and *Plac8* within the macrophages present pre- and post-birth. (E) Multiplexed *in situ* hybridization to detect gene expression of *Cd68* (yellow), *Dab2* (red), and *Plac8* (green) in lung tissue from mice at E18.5 and P7. Calibration bar=20μm.

The Mac I-V clusters broadly separated into two groups based upon expression of the disabled 2 gene (*Dab2)*, which regulates macrophage polarization^23^ and the placenta-specific 8 gen*e (Plac8)*, which is related to bacterial immunity^24^. Mac I-IV cells expressed *Dab2* but not *Plac8* while Mac V showed the opposite pattern (Figure 2D). We confirmed by multiplexed FISH that all *Cd68+* cells in the lung expressed either *Dab2* or *Plac8* at both E18.5 and P7 (Figure 2E). *Dab2* and *Plac8* expression was not consistent with previously reported markers of macrophage lineages derived from yolk sac and fetal liver^25, 26^ (Supplemental Figure 2A).

### Embryonic macrophages are proliferative and encircle developing vessels prior to birth

Mac I cells are the predominant immune population at E18.5, hence we aimed to understand their function and localization. Differentially expressed genes (DEGs) in Mac I included the proliferation markers *Mki67* and *Mcm5* (Figure 3A). Across macrophages and monocytes, proliferation decreased from 60% of cells at E18.5 to only 10% by P21. Most proliferating cells were Mac I prior to birth and distributed across Mac II-V after birth (Supplemental Figure 2B). These data are consistent with prior reports of “bursts” of proliferation after recruitment of macrophages into embryonic tissues, followed by low-level self-renewal by adulthood^27^.

**Figure 3:**
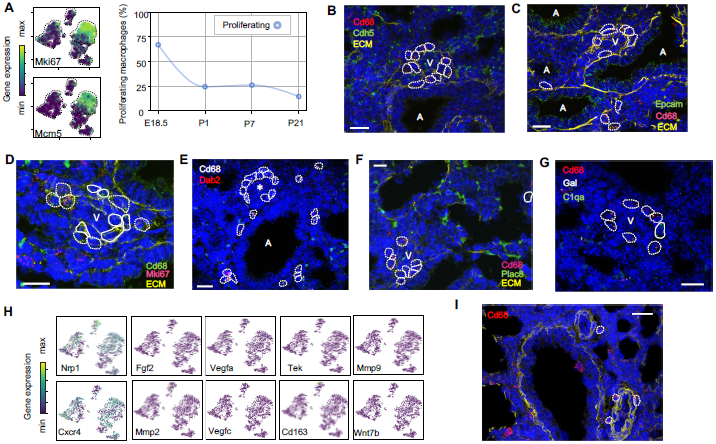
Embryonic macrophages encircle developing blood vessels prior to birth. (A) t-SNE plots depicting expression of *Mki67* and *Mcm5* in the macrophage clusters with low expression in purple and high expression in yellow, with quantification of proliferating macrophages at each timepoint. *In situ* hybridization at E18.5 to detect: (B) *Cd68* (red) and *Cdh5* (green) demonstrating circles of macrophages around small vessels; (C) *Epcam* (green), *Cd68* (red), and extracellular matrix (ECM, yellow), with white dotted circles identifying *Cd68+* cells; (D) *Mki67* (red), *Cd68* (green), and ECM (yellow), with white dotted circles identifying *Cd68*^+^*Mki67*^+^ cells, and solid circles identify *Cd68*^+^ *Mki67*^-^ cells; (E) *Cd68* (white) and *Dab2* (red) with white dotted circles identifying *Cd68*^+^*Dab2*^+^ macrophages; (F) *Plac8* (green), *Cd68* (red), and ECM (yellow), with white dotted circles identifying *Cd68*^*+*^*Plac8*^*-*^ cells and solid circles *Cd68*^*+*^*Plac8*^*+*^ cells; (G) *Cd68, C1qa*, and *Gal*. (H) t-SNE plots of genes previously associated with a perivascular macrophage phenotype. (I) *In situ* hybridization of lung at P1 to detect *Cd68* (red) and ECM (yellow), with white dotted circles identifying isolated macrophages around blood vessels.

Localization of Mac I cells within the E18.5 lung revealed *Cd68+* cells scattered throughout the lung parenchyma but also, surprisingly, forming almost complete rings around blood vessels of 20-30 μm in diameter found adjacent to large, conducting airways (Figure 3B). In contrast, Mac I cells were not found encircling small airways (Figure 3C). Many of the vessel-surrounding macrophages expressed *Mki67* (Figure 3D) and *Dab2* (Figure 3E) but not *Plac8* (Figure 3F). Given that *Dab2*+ cells at E18.5 include Mac IV cells, we aimed to distinguish these from the Mac I cells by detecting the expression of *Gal* and *C1qa*, which were determined to be specific markers by scRNA-Seq (see below). These studies demonstrated that the majority of *Cd68*+ macrophages surrounding the vessels were *Gal*+ with an occasional *C1qa*+ cell, indicating a predominance of Mac I and a small number of Mac IV cells comprise the perivascular macrophage population (Figure 3G). We then asked whether any of our Mac I-V clusters are related to previously reported perivascular macrophages that promote vascular remodeling in the developing hindbrain and retina^28^. Mac I cluster expressed *Cxcr4* and *Nrp1* but failed to express many other genes characteristic of these previously reported macrophages, suggesting a distinct phenotype (Figure 3H). Moreover, no concentric perivascular macrophages were observed at P1, consistent with a function specific to Mac I prior to birth (Figure 3I). Taken together, these data suggest that within the embryonic lung, Mac I macrophages are highly proliferative and localize to small vessels, suggesting a potential role in pulmonary vascular growth or remodeling.

### Distinct transcriptional profiles and spatial distribution suggest specific physiologic functions for discrete macrophage populations

Macrophage and monocyte heterogeneity increased rapidly after birth. To characterize this rising diversity we computed DEGs for each of the Mac I-V clusters (Figure 4A and Supplemental Tables 1-5). Beyond proliferation, the embryonic cluster Mac I expressed genes associated with glycolysis, reflective of the hypoxic fetal environment compared to postnatal air-breathing life. Mac I-specific DEGs also included *Crispld2*, a glucocorticoid-regulated gene previously thought to be restricted primarily to the lung mesenchyme that promotes lung branching^29^. *Crispld2* haploinsufficient mice exhibit impaired alveolarization and disorganized elastin deposition^30^. Mac I cells also expressed *Spint1*, encoding hepatocyte growth factor activator inhibitor type 1 (HAI-1), a membrane bound serine proteinase inhibitor and regulator of angiogenesis (Figure 4A and Supplemental Table 1). Loss of *Spint1*, results in a complete failure of placental vascularization and embryonic lethality at E10 that appears to result from a loss of basement membrane integrity^34^. The most specific marker for Mac I was *Gal*, encoding galanin, a regulatory peptide that harbors both pro- and anti-inflammatory functions^35^, promotes an anti-thrombotic phenotype in endocardial EC^36^, and regulates growth and self-renewal of embryonic stem cells^37^. Galanin also inhibits inflammatory and histamine-induced vascular permeability in a number of experimental models ^38-40^, and functions as a vasoconstrictor, limiting blood flow in the cutaneous microcirculation^41^. Localization identified Mac I cells throughout the lung parenchyma in addition to those found encircling small vessels (Figure 4B).

**Figure 4:**
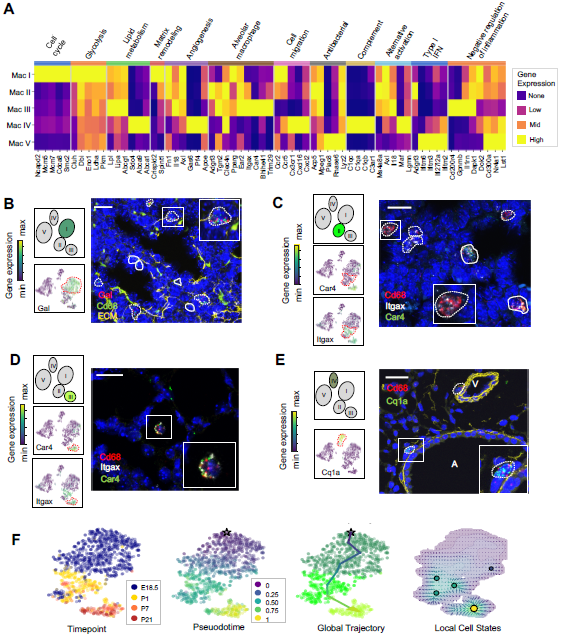
Distinct transcriptional profiles and spatial distribution suggest specific physiologic functions for discrete macrophage populations. (A) Heatmap of select differentially expressed genes within enriched pathways illustrated. (B) t-SNE plots demonstrating high expression of Gal in Mac I cells, and *in situ* hybridization at E18.5 to detect Mac I cells that co-express *Gal* (red) and *Cd68* (green). (C) t-SNE plots demonstrating high expression of *Itgax* but not *Car4* in Mac II cells, and *in situ* hybridization to detect Mac II cells at P1 expressing *Itgax*, and *Cd68* but not *Car4* (dotted line), and additional Mac III cells in the same image co-expressing *Itgax, Cd68*, and *Car4* (solid line). (D) t-SNE plots demonstrating high expression of *Itgax* and *Car4* in Mac III cells, and *in situ* hybridization detecting Mac III cells at P7 expressing *Itgax, Car4 and Cd68* now located within alveoli. (E) t-SNE plot demonstrating high expression of *C1qa* in Mac IV cells, and *in situ* hybridization to detect Mac IV cells expressing *C1qa (green)* and *Cd68(red)* at P7, with ECM marked in yellow, localizing Mac IV cells abutting vessels and large airways. Calibration bar=20 mm for all panels. (F) t-SNE plot showing a developmental gradient across Mac I-III. Pseudotime ordering of the cells identified a global trajectory from the starting cell (star) in Mac I to Mac III, and local cell states revealing multiple areas of local attraction within the Mac II and Mac III clusters.

The Mac II cluster rapidly appeared after birth and expressed a gene signature suggesting a putative role in immune regulation and tissue remodeling. Mac II cells shared expression of the chemokine receptors *Ccr2* and *Ccr5* with Mac I, molecules important for immune cell migration and localization. Also similar to Mac I and III, Mac II cells expressed genes associated with matrix remodeling and angiogenesis (*Fn1* and *Axl*)^32, 33^ (Figure 4A and Supplemental Table 2). The Mac II cluster also shared genes with Mac III important for regulating inflammation, including genes that promote intracellular killing of pathogens (*Acp5* and *Mpeg1*)^34^, but also genes that suppress inflammation (*Il1rn and Dapk1*)^35, 36^. A subpopulation within Mac II shared high expression of major histocompatibility complex (MHC) class II genes (*H2-Ab1, H2-Eb1, Cd74*) with a subset of Mac IV (Supplemental Fig. 3) suggesting a role in antigen presentation. Overall, the transient presence of Mac II together with its transcriptional signature suggests a dual role in tissue remodeling and fine tuning of immune response required immediately after birth. Studies to localize *Itgax*+*Car4*-Mac II cells *in situ* at P1 identified them in the distal lung parenchyma, mixed with scattered Mac III cells which were located closer to the alveolar wall (Figure 4C and Supplemental Figure 4A).

The Mac III cluster uniquely expressed alveolar macrophage signature genes (*Car4, Bhlhe41, Trim29*) (Figure 4A, Supplemental Table 3), and genes that constrain inflammation (*Cd200r4, Gpnmb, Il1rn*)^37, 38^. Mac III cells also expressed genes *Lpl, Lipa*, and *Abcg1*, indicative of their essential role in surfactant catabolism^39, 40^. *In situ* imaging of *Itgax+ Car4+* cells demonstrated that they move from the distal lung parenchyma at P1 to the alveolar lumen by P7 (Figure 4D), confirming their identification as alveolar macrophages (AMs).

Mac IV uniquely expressed numerous proinflammatory genes^41^ (Figure 4A), in contrast to the balanced inflammatory signature of Mac II and AMs. These included genes in the classical complement pathway (*C1qa, C1qb, C1qc, C3ar1*) and CCR2 ligand *Ccl12* (Supplemental Table 4), suggesting a role in the localization of CCR2 expressing monocytes^42^. The Mac IV cells also expressed *Cxcl16*, an IFNγ regulated chemokine that promotes T cell recruitment through CXCR6^43^. Mac IV also highly expressed *Mrc1* (CD206) (Supplemental Fig. 3A). However, within the Mac IV cluster there were a group of cells with lower *Mrc1* and high MHC II gene expression (*H2-Ab1, H2-Eb1*, and *Cd74*). These transcriptional differences within Mac IV are similar to two recently reported interstitial macrophages (IMs) in adult lung that can be differentiated by expression of Mrc1 and MHC II genes^44^. However, the other genes reported to distinguish those two clusters (e.g *Cd68, Cx3cr1, Mertk, Cclr2*) were diffusely expressed throughout the Mac IV cluster (Supplemental Fig. 3B). *In situ* imaging to localize *C1qa*+*Cd68*+ cells in the postnatal lung found Mac IV cells remained abutting small blood vessels as well as the abluminal side of large airways (Figure 4E). The characteristic location and the expression of numerous genes important for leukocyte recruitment and pathogen defense suggest that Mac IV may serve as patrollers, playing a key role in immune-surveillance, innate pathogen defense, and antigen presentation.

The transient presence of the Mac II cells, and significant overlap with the transcriptomes of Mac I, and III, suggested that Mac II may represent an intermediate population. To determine if cells within these clusters were undergoing gradual transcriptional shifts across time, we performed pseudotime analysis on cells from clusters Mac I, II, and III. Ordering of the cells within the Mac clusters I-III by pseudotime (Figure 4F) defined a global trajectory from the Mac I to the Mac III cluster, indicating a correspondence between pseudo- and real time during perinatal development. This global trajectory notwithstanding, there were multiple local attractor states within the Mac II cluster, suggesting that cells gradually shift from Mac I to Mac II, and subsequently remain in that phenotypic state for some time before further committing to a specific fate. Interestingly, one of these local attractor states corresponded with the subpopulation expressing MHC class II genes also found in a subset of Mac IV cells. Taken together, our data suggest that the Mac II cells derive from Mac I and represents a transitional population that may serve a temporal-specific function and later differentiates into Mac III cells (i.e. alveolar macrophages) and potentially the antigen-presenting subset of Mac IV cells.

### Mac V monocytes are characterized by developmental gene expression gradients

The Mac V cluster was characterized by expression of *Plac8*^*4*^ (Figure 1E), a gene expressed by a recently identified population of “CD64+ CD16.2+ non-classical monocytes” in the adult mouse lung^44^, suggesting a monocytic phenotype. Mac V monocytes also expressed a panel of unique transcripts induced by type I interferon (IFN), important for modulating host responses to viral pathogens (*Ifitm2, Ifitm3, Ifitm6, Ifi27l2a*)^46^ (Figure 5A), as well as genes that constrain inflammation (*Cd300a, Nr4a1, Lst1*) (Figure 4A and Supplemental Table 5). This dual immune signature was similar to that seen in Mac II and III, emphasizing the importance of a finely tuned inflammatory response in the developing lung.

**Figure 5:**
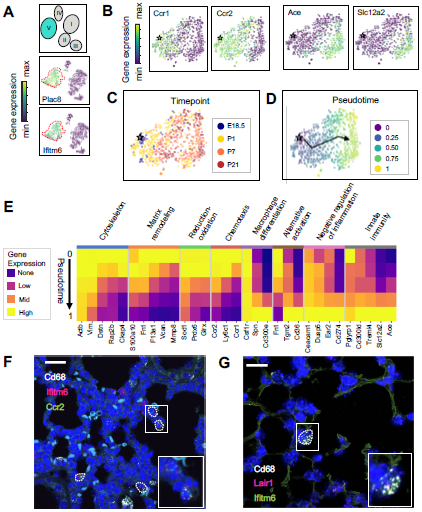
The Mac V cluster harbors distinct subpopulations with developmentally regulated gene expression patterns. (A) t-SNE plots demonstrating high expression of *Plac8 and Ifitm6* in Mac V cells. (B) t-SNE plots of *Ccr1, Ccr2, Ace*, and *Slc12a2* suggesting the presence of two transcriptionally distinct populations within the Mac V cluster. (C) t-SNE demonstrating a developmental gradient within the Mac V sub cluster. (D) Pseudotime analysis with the star indicating the starting point and the arrow denoting the trajectory across pseudotime. (E) Heatmap of differentially expressed genes within enriched pathways across pseudotime. (F) *In situ* hybridization of *Cd68 (white), Ifitm6 (red), and Ccr2 (green)*, and to detect the “early” Mac V subcluster at P1. (G) *In situ* hybridization of *Cd68 (white), Lair1 (red), and Ifitm6 (green)*, to detect the “late” Mac V subcluster at P21. Calibration bar=20mm for all panels.

Within Mac V we found a gradient of cell states between two distinct phenotypes, with some genes (e.g. *Ccr1* and *Ccr2)* expressed at one end and other genes (e.g. *Ace* and *Slc12a)* expressed at the other end of the spectrum (Figure 5B), with a clear correspondence to real developmental time (Figure 5C). Pseudotime analysis indicated that cells with an early phenotype upregulated *Ly6C*^*4*^, a gene expressed by classical monocytes, and genes associated with the cytoskeleton (*Actb, Vim*)^48, 49^, matrix remodeling (*Fn1, F13a1, Vcan*)^50, 51^, and reduction-oxidation (*Sod1, Prdx6*)^52, 53^ in keeping with the marked physiological changes and rapid remodeling occurring in the lung during the fetal-neonatal transition (Figure 5D and E). Over both pseudo- and real time, this gene expression pattern evolved into an immunomodulatory signature, reflected by the up-regulation of genes associated with macrophage differentiation (*Csf1, Spn*)^54^, macrophage polarization (*Tgm2, Cd36*)^55^, negative regulation of inflammation (*Ceacam1, Ear2, Lair1*)^56^, and innate immunity (*Cd300ld, Treml4*)^57^. These data indicate that only the late Mac V cells are similar to the CD64+ CD16.2+ non-classical monocytes reported by Schyns et. al.^44^, while the early Mac V cells represent an additional source of heterogeneity unique to the perinatal lung.

Spatial localization of the early and late populations by detecting either *Ccr2* or *Lair1* in combination with *Ifitm6* and *Cd68*, allowed the identification of early Mac V cells within the distal lung parenchyma (Figure 5F, Supplemental Figure 4B). Of note, these *Cd68*+ *Ifitm6*+ *Ccr2*+ cells were reliably found either along the secondary crests or lining the developing alveoli such that one boundary of the cell was always in contact with the airspace. Despite significant changes in the gene expression, *Cd68*+ *Ifitm6+Lair1+* were similarly found in the distal lung along the alveolar walls (Figure 5G). Taken together, these data demonstrate that within the Mac V monocytic phenotype there are functionally distinct, developmentally dynamic cell states that transition during and after birth from tissue remodeling and regulation of immune cell chemotaxis to immunomodulation and pathogen defense.

### Variations in macrophage Fc receptor expression

Fc receptors serve as an important link between cellular and humoral immunity by bridging antibody specificity to effector cells^58^ and are therefore a key axis of functional heterogeneity within macrophages and monocytes. We evaluated their expression across Mac I-V and found specific patterns (Supplemental Figure 5). Both *Fcgr3* (encoding FcγRIII) and *Fcer1g* (FcεRIγ) were widely expressed by all five clusters, while *Fcer1a* and *Fcer2a* were not expressed by any cluster. Expression of *Fcgr1*, a high affinity receptor for IgG important for the endocytosis of soluble IgG, phagocytosis of immune complexes, and delivery of immune complexes to APC^59^ was highly expressed by Mac I and Mac IV, and in the early subcluster of Mac V. In contrast, *Fcgr4*, encoding an Fc receptor able to bind IgE that promotes allergic lung inflammation was highly expressed by the late sub-cluster of Mac V, in agreement with data from adult mice^44, 60, 61^. *Fcgrt*, encoding the neonatal Fc receptor (FcRn), an Fc receptor with a key role in IgG recycling, was highest in Mac IV cells.

### Dendritic cell subtypes and granulocytes are primed for rapid pathogen response

Dendritic cells (DCs) play multiple roles in the immune system including antigen presentation and regulation of tolerance and can be distinguished from other mononuclear phagocytes by the expression of *Zbtb46*^*6*^ and *Flt3*^*6*^. We found three clusters of cells, DC I-III, (Figure 6A and Supplemental Tables 6-8) expressing *Zbtb46* and *Flt3* (Supplemental Figure 6A). DC I cells expressed *Itgae* or CD103 (Figure 6B), which promotes antiviral immunity and may confer the ability for antigen cross-presentation^64^. DC II expressed *Cd209a* or DC-SIGN, a gene found in monocyte-derived inflammatory DC exposed to lipopolysaccharide^65^. DC III expressed melanoregulin (*Mreg*), a modulator of lysosomal hydrolase maturation^66^ (Figure 6B). *Mreg* had not been previously identified as a marker for DC subsets, hence we examined other genes expressed by DC III and identified *Cacnb3*, a voltage dependent Ca^2+^ channel found in stimulated Langerhans cells^67^; *Fscn1*, which contributes to dendrite formation in maturing DC^68^; *Ccl5*, an important chemoattractant for DC and T cells; and *Ccr7*, a chemokine receptor associated with trafficking to the draining lymph node^69^ (Supplemental Figure 6B and Supplemental Tables 6-8).

**Figure 6:**
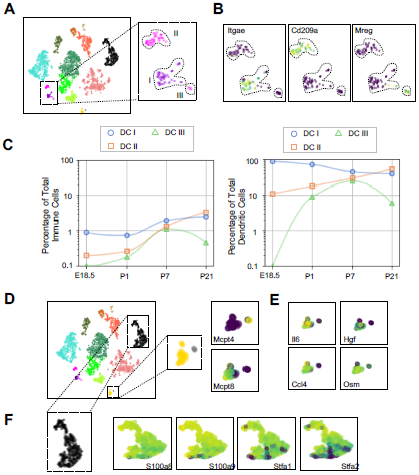
Multiple dendritic cell populations and lung granulocytes are primed for rapid pathogen response. (A) Colored schematic and lung immune cell clustering demonstrating three separate clusters of DCs. (B) tSNE plots of genes discriminating the three DC subclusters including *Itgae (*DCI*), Cd209a (*DCII*) and Mreg (*DCIII). (C) Quantification of specific DC subpopulations relative to total immune cells (left) or total DC (right). (D) Colored schematic and lung immune cell clustering demonstrating the basophil, mast cell and neutrophil clusters, with high magnification of basophil and mast cell clusters and t-SNE plots of *Mcpt4* and *Mcpt8*. (E) t-SNE plots of *Il6, Hgf, Ccl4*, and *Osm* in the basophil cluster. (F) High magnification of the neutrophil cluster with t-SNE plots of neutrophil specific genes *S100a8, S100a9, Stfa1*, and *Stfa2*.

Quantification of the relative abundances identified DC I as the predominant population in the embryonic lung. DC I persisted postnatally to comprise between 1-2% of total lung immune cells (Figure 6C). DC II was present at low frequency during the first week of life and increased in abundance by P21. DC III was undetectable before birth and became more abundant postnatally. Taken together, these data suggest DC I comprises migratory DCs, DC II cells are related to monocyte-derived DCs, and DCIII is a minority subtype of mature DCs.

Mast cells and basophils are similar in development and function and serve as fast responders to specific immune challenges^70^. Two immune clusters highly expressed *Cpa3* that could be further distinguished as mast cells and basophils by the expression of *Mcpt4* and *Mcpt8*, respectively^71, 72^ (Figure 6D). Mast cells expressed *Tpsb2*, which is secreted upon bacterial challenge^73^, and the peptidases chymase (*Cma1*) and tryptase (*Tspab1*) (Supplemental Tables 9). Lung resident basophils express a unique signature distinct from circulating basophils and play a key role in promoting AM differentiation^74^. Our basophils generally shared expression of many genes with lung resident basophils, including *Il6, Hgf, Ccl4* and *Osm* (Figure 6E, Supplemental Table 10). We also identified a neutrophil cluster distinguished by expression of *S100a8* and *S100a9*, which are released during inflammation^75^, and *Stfa1* and *Stfa2*, cysteine proteinase inhibitors important for antigen presentation^76^ (Figure 6F and Supplemental Table 11). Our data agree with prior work demonstrating that lung resident basophils express signaling molecules important for interaction with neighboring cells^74^, and that mast cells and neutrophils are primed for rapid innate immune responses upon pathogen challenge.

### Naïve lymphocytes populate the lung at birth

Lymphocytes including ILC2s (expressing *Areg*), NK cells (*Gzma*), B cells (*Ms4a1)* and T cells (*Cde3)* were present in the lung at low frequencies (approximately 2% of immune cells) prior to birth and increased in frequency after birth. By P21 lymphocytes comprised 60% of total immune cells (Figure 1F and G), with B cells representing 30% and T cells 15% of the total (Figure 7A).

**Figure 7:**
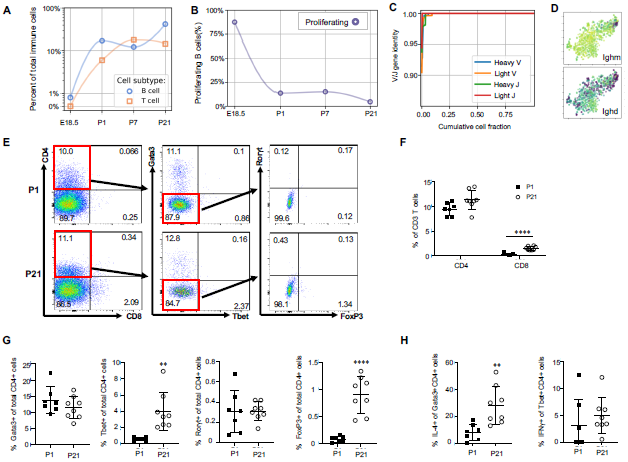
Lymphocytes populate the lung at birth but remain naïve during the first three weeks of life. (A) Quantification of the abundance of B and T cells at each developmental timepoint expressed on a log10 scale as percentage of total immune cells. (B) Quantification of the percentage of proliferating B cells at each developmental timepoint. (C) B cell heavy and light variable (V) and joining (J) gene identity and their cumulative cell fraction. (D) t-SNE plot of *Ighm* and *Ighd*. (E-H) At P1 (n=7 mice) and P21 (n=8 mice), lungs were processed to a single-cell suspension, and flow cytometry was used to assess frequencies of (F) CD4+CD3+ and CD8+CD3+ T cells, (G) Gata3+, Tbet+, Rorγt+ and Foxp3+ CD4+ T cells, and (H) IL-4-producing Gata3+CD4+ T cells or IFNγ-producing Tbet+CD4+ T cells. Data shown as mean ± SD, \*\**P* < 0.01, \*\*\*\**P* < 0.00001 by Student’s *t* test.

B cells in the embryonic lung were rare and the majority expressed proliferation markers. After birth, B cell abundance increased, but the proliferating fraction decreased (Figure 7B). The cumulative distribution of V and J loci germline similarity revealed no somatic hypermutation (Figure 7C) and most B cells expressed *Ighm* and *Ighd* indicative of an IgM isotype (Figure 7D). Few B cells expressed activation markers (e.g. *Aicda, Tbx21, Prdm1*, and *Ebi3*, Supplemental Figure 7B)^77-80^, suggesting that most postnatal B cells remain naïve through late alveolarization. To test the clonality of the B cell repertoire at birth, we assembled the heavy and light chain loci and performed t-SNE on a feature-selected transcriptome limited to over-dispersed genes in B cells. Proliferating cells clustered together but candidate clonal families did not, suggesting primarily homeostatic B cell proliferation rather than clonal expansion^81^.

The majority of T cells expressed *Trac*, suggesting αβ identity, with a few *Trac-* cells expressed *Tcrg-C4*, suggesting γδ T cell identity (Supplemental Figure 7C). T cell receptor diversity showed no sign of clonal expansion (data not shown). Outside the thymus, αβ T cells are usually either CD4+ (with subgroups Tbet+ or T helper (Th) 1, Gata3+ or Th2, and FoxP3+ or Th17) or CD8+, however we characterized T cell heterogeneity in neonatal lungs by flow cytometry and found that 85-90% of CD3+ cells were CD4- CD8- at both P1 and P21 (Figure 7E), confirming an earlier report suggesting this is a neonatal-specific phenotype. mRNA expression analysis qualitatively confirmed this finding (Supplemental Figure 7D). The frequency of total T cells was similar at P1 and P21 (Supplemental Figure 7E), however several T cell subsets (CD8^+^, CD4+ Th1, and Treg cells) increased by P21 while other subsets remained constant (Figure 7F and G). In response to stimulation with phorbol myristate acetate (PMA) and ionomycin, a greater number of CD4^+^ Th2 cells at P21 produced IL-4 as compared to P1 (28.3 ± 19.7 vs. 8.6 ± 5.7, P=0.0047) (Figure 7H). Few CD4+ Th1, Treg, and Th17 cells produced IFNγ, IL-10, IL-17 upon stimulation, at both P1 and P21 (Figure 7H, Supplemental Figure 7F).

## Discussion

At birth, the lung undergoes marked physiological changes as it transitions from a fluid-filled, hypoxic environment to an air-filled, oxygen-rich environment. How these changes affect immune populations during this transition and the ensuing period of rapid postnatal lung growth remains unclear. Our study demonstrates a rapid increase in immune cell heterogeneity, especially within macrophages and monocytes. We identified five macrophage subpopulations, each expressing a specific gene signature, spatial localization, and putative functions. Mac I, the predominant immune cell present just before birth, were highly proliferative, enriched for tissue remodeling and angiogenesis genes, and completely encircled small blood vessels, suggesting a previously unrecognized role for lung macrophages in modulating lung vascular growth or remodeling during development. During the first week of life a transitory population (Mac II) emerged from Mac I and later disappeared, transitioning into either an alveolar (Mac III) or Mac IV macrophage phenotype. One macrophage population (Mac IV) expressed complement proteins and other antibacterial molecules. Another interstitial population (Mac V) expressed antiviral molecules and spanned a gradient between two extreme phenotypes, one that expressed high levels of homeostatic genes during early postnatal development, and a second with immunomodulatory function that resembles previously reported nonclassical monocytes. Lymphocytes increased in abundance from almost zero before birth to more than half of lung immune cells by P21, but maintained a naive phenotype skewed toward type II immunity and with predominantly Cd4- Cd8- T cells.

This comprehensive study has far-reaching implications for lung biology. Resident tissue macrophage populations are established during development, wherein progenitors undergo differentiation guided by the tissue-specific microenvironment^82^. However, definitive data regarding the full complexity of lung resident macrophages and monocytes, their specific roles and functions, and how they change across development remain elusive. Although the advent of single cell transcriptomics has provided increased resolution to detect previously unrecognized immune cell populations, consensus regarding the diversity and function of lung resident macrophages has not been achieved. Cohen et al. recently performed single cell transcriptomics of the developing mouse lung from E12.5 until P7^74^, and identified a total of three macrophage populations, and one population of resident monocytes, with alveolar macrophages representing the sole macrophage population present in the lung after P7. In contrast, Schyns et al. identified two distinct interstitial macrophages in the adult lung, and a population of nonclassical monocytes in addition to alveolar macrophages^44^. Although our results are more consistent with the report of Schyns et al, there are a number of key differences. First, the total heterogeneity in the perinatal lung far exceeds the adult lung, with the presence of two unique macrophage clusters (Mac I and Mac II and a unique monocyte derived cluster (early Mac V). Second, both the Mac IV and Mac V cluster harbor significant internal heterogeneity (in the case of Mac V, corresponding to developmental time) that cannot be easily split into transcriptionally distinct “subclusters”. The cells within Mac IV appear similar to the CD206+ and CD206-macrophage populations reported by Schyns et al. Although in the Schyns report those populations were reported to be distinct clusters, there was significant overlap in gene expression between the two, more consistent with our data suggesting these are not separate populations but rather a phenotypic continuum. *In situ* validation confirmed the presence of all five subpopulations and localized each to defined locations in the lung including the alveolar lumen, around vessels and airways, or within the distal lung interstitium. A greater understanding of macrophage and monocyte function at birth provides an essential framework for interpreting how lung injury and developmental defects alter specific immune subpopulations and eventually influence lung growth and development.

Another key finding in our study was the unexpected presence of embryonic lung macrophages encircling small blood vessels. Vascular growth is a key driver of distal lung growth during the late saccular and alveolar stages of development^83^. Macrophages support angiogenesis in other organs, promoting blood vessel formation or expansion, providing survival and migratory cues to endothelial cells, and facilitating bridging of vascular sprouts^84^. In the developing hindbrain, macrophages are in close contact with endothelial cells, serving to promote vascular anastomosis^85^. Similarly, in the developing retinal vasculature, microglia connect adjacent endothelial tips cells to increase vascular plexus complexity^9^. These embryonic bridging macrophages secrete numerous genes shared by the perivascular macrophages that drive tumor angiogenesis including the angiopoietin receptor, *Tek*, the VEGF co-receptor *Nrp1*, growth factors (*Fgf2, Pgf*) and MMPs (*Mmp2, Mmp9*). Although the perivascular macrophages we observed in the embryonic lung expressed low levels of *Nrp1*, they appear distinct from the macrophages that influence retinal and hindbrain angiogenesis, expressing a unique set of ECM remodeling and angiogenic genes, including genes that may modulate vascular tone and permeability. The distinctive location of these macrophages and their gene signature imply a role in vascular development. Furthermore, these encircling macrophages disappeared after birth, suggesting a function temporally restricted to prenatal development. Future studies to selectively target this subpopulation will be required to further establish their function and to delineate the signals responsible for the cessation of the macrophage-vascular interaction after birth.

During embryonic development, the lung is populated by separate populations of erythroid-myeloid progenitors originating from the yolk-sac and fetal liver, prior to the emergence of circulating monocytes and hematopoietic stem cells. Some existing data suggest that the early yolk sac derived macrophages are eventually entirely replaced by fetal liver derived macrophages capable of self-renewal^86^. In our study, we observed a broad division of the five macrophage populations based upon expression of *Dab2* and *Plac8*, evident both before and after birth. However, expression of genes that characterize yolk sac- and fetal liver-derived macrophages at earlier stages of development were dispersed among all five populations^25^. These data are consistent with prior work suggesting that imprinting from signals in the tissue microenvironment is the dominant factor regulating macrophage phenotype^87^. The additional contribution of bone marrow derived monocytes to replenish lung macrophages under homeostatic conditions also remains debated^88^. High expression of *Ly6C* and *Ccr2* observed in early Mac V cells is reminiscent of the infiltrating monocytes that continuously replenish intestinal macrophages, indicating that both early and late Mac V cells are monocytes or monocyte-derived^89^. Future experiments exploiting lineage tracing technologies at perinatal time points are warranted to determine the amount of extravasating blood-derived versus self-sustaining tissue-resident monocytes at this crucial time.

Our data also revealed three distinct dendritic cell populations with transcriptional signatures indicative of migratory, inflammatory, and a mature dendritic cell phenotypes, respectively. Moreover, we identified distinct (*Itgae* and *Cd209a*) and novel *(Mreg)* markers superior to classical markers (*Zbtb46* and *Flt3*) to distinguish dendritic subpopulations in the postnatal lung. Functionally, both *Itgae* and *Cd209a/*DC-SIGN can induce T cell immunity^90, 91^, suggesting that lung DCs immediately post-birth should be able to cause an effective adaptive immune response.

Despite the apparent signaling readiness of antigen-presenting DCs, T and B cell compartments showed an overall naive and rarely proliferative phenotype, lacking any clonal structure and with most αβ T cells double negative (DN) for both Cd4 and Cd8, key signaling components for cell-mediated immunity. Though consistent with prior evidence showing a high proportion of pulmonary lymphocytes with unconventional phenotypes^92^, the observation of widespread DN T cells points to a yet undetermined function, perhaps related to specific immunoregulatory functions during infectious disease^93^.

In summary, these data highlight the marked increase in immune cell diversity after birth, with a developmental plasticity that provides distinct immune populations to fill specific roles in tissue and vascular remodeling, immunoregulation, and bacterial and viral pathogen defense. Injuries to the developing, immature lung can have profoundly untoward and life-long consequences as a significant component of lung parenchymal and vascular development occurs during late pregnancy and the first few years of postnatal life. Many of these injurious stimuli including acute infection, hyperoxia, and corticosteroids are known to have significant effects on immune cell phenotype and function. Therefore, our data provide a detailed framework that enables a more complete understanding of how disruptions of immune cell phenotype may contribute to altered lung development, both through the induction of pathologic, pro-inflammatory signaling as well as the suppression of essential homeostatic functions. Further, a deep understanding of the diversity of immune cell functions during this important window of postnatal development, and how specific immune cell phenotypes are regulated could allow for the application of immunomodulatory therapies as a novel strategy to preserve or enhance lung development in infants and young children.

## Acknowledgements

We thank Sai Saroja Kolluru (Stanford University) for assistance with library submission to the Chan Zuckerberg Biohub, Yuan Xue (Stanford University) for assistance with the initial single cell RNA-seq data acquisition, Astrid Gillich for technical support with the RNAscope experiments, and Maya Kumar for providing hydrazide. We thank Henry Hampton for his input and fruitful discussions. We also thank the Stanford Shared FACS Facility, Lisa Nichols, Meredith Weglarz, and Tim Knaak for assistance with the flow cytometry instrumentation and antibody panel design. Flow cytometry data was collected on an instrument in the Stanford Shared FACS Facility obtained using NIH S10 Shared Instrument Grant (S10RR027431-01). This work was supported by National Institutes of Health grants HL122918 (CMA), HD092316 (CMA, DNC), the Stanford Maternal Child Health Institute Tashia and John Morgridge Faculty Scholar Award (CMA), the Stanford Center of Excellence in Pulmonary Biology (DNC), Bill and Melinda Gates Foundation (SRQ), and the Chan Zuckerberg Biohub (DNC. and SRQ). MAS is supported by the NSF-GRFP.

## Author Contributions

R.D.-G., C.M.A., F.Z., X.C., S.R.Q., and D.N.C. designed the experiments, interpreted the data, and wrote or edited the manuscript. R.D.-G., X.C., M.L., and R.C.J. performed the experiments. R.D.G., X.C., and F.Z. prepared the sequencing libraries. F.Z. analyzed the transcriptomic data. M.A.S. assembled the B and T cell repertoires. All authors edited and approved the final version of the manuscript.

## Methods

### Mouse lung cell isolation

C57BL/6 mice were obtained from Charles River Laboratories. For studies using E18.5, P1, and P7 murine lungs, pregnant dams were purchased, and pups aged prior to lung isolation. At E18.5, dam was asphyxiated with CO2 and pups extracted. At P1, P7, and P21 pups were euthanized with euthanasia solution (Vedco Inc.). Genetic sex of mice at developmental stages E18.5 and P1 was determined by performing PCR amplification of the Y chromosome gene Sry. P7 and P21 mice were sexed through identification of a pigment spot on the scrotum of male mice^94^. For all timepoints, except E18.5, the pulmonary circulation was perfused with ice cold heparin in 1x PBS until the circulation was cleared of blood. Lungs were minced and digested with Liberase (Sigma Aldrich) in RPMI for 15 (E18.5, P1, and P7) or 30 (P21) minutes at 37C, 200 rpm. Lungs were manually triturated and 5% fetal bovine serum (FBS) in 1x PBS was used to quench liberase solution. Red blood cells were lysed with 1x RBC lysis buffer (Invitrogen) as indicated by the manufacturer and total lung cells counted on Biorad cell counter (BioRad).

### Immunostaining and fluorescence-activated cell sorting (FACS) of single cells

Lungs were plated at 1×10^6^ cells per well and stained with Fc block (CD16/32, 1:100, Tonbo Biosciences) for 30 min on ice. Cells were surface stained with the endothelial marker CD31 (1:100, clone: MEC3.1, eBiosciences), epithelial marker Epcam (1:100, clone: CD326, eBiosciences), and immune marker CD45 (1:100, clone: F11, eBiosciences) for 30 min on ice. The live/dead dye, Sytox Blue (Invitrogen), was added to cells and incubated for 3 min prior to sorting into 384-well plates (Bio-Rad Laboratories, Inc) using the Sony LE-SH800 cell sorter (Sony Biotechnology Inc). FACS sorts were performed with a 100 μm sorting chip (Catalog number: LE-C3110). Prior to cell sorting, the cell sorter and chip were calibrated with SH800 setup beads. Droplet targeting into the middle of four corner and center wells of 384-well plates was manually calibrated. Single color controls were used to perform fluorescence compensation and generate sorting gates. The 384-well plates pre-loaded with lysis buffer (Triton X-100 solution, dNTP, poly dT, RNase inhibitor, ERCC, and Triton X-100) were loaded onto the Sony SH800 for gated single cell capture using the ultra-purity mode. Following completion of sorting, 384-well plates containing single cells were spun down, immediately placed on dry ice, and stored at −80C.

### cDNA library generation using Smart-Seq2

Complementary DNA from sorted cells was reverse transcribed and amplified using the Smart-Seq2 protocol on 384-well plates as previously described^20, 95^. Concentration of cDNA was quantified using picogreen (Life technology corp.) to ensure adequate cDNA amplification. In preparation for library generation, cDNA was normalized to 0.4 ng/uL. Tagmentation and barcoding of cDNA was prepared using in-house Tn5 transposase and custom, double barcoded indices^96^. Library fragment concentration and purity were quantified by Agilent bioanalyzer. Libraries were pooled and sequenced on Illumina NovaSeq 6000 with 2×100 base kits and at a depth of around 1 million read pairs per cell.

### Data analysis and availability

Sequencing reads were mapped against the mouse genome (GRCm38) using <u>STAR aligner^97^</u> and gene were counted using <u>HTSeq^98^</u>. FZ has been the main maintainer of HTSeq for several years. To coordinate mapping and counting on Stanford’s high-performance computing cluster, <u>snakemake</u> was used^99^. Gene expression count tables were converted into loom objects (https://linnarssonlab.org/loompy/) and cells with less than 100,000 uniquely mapped counts were discarded. Counts for the remaining cells were normalized to counts per million reads. For t-distributed stochastic embedding (t-SNE)^22^, 500 features were selected that had a high Fano factor in most mice, and the restricted count matrix was log-transformed with a pseudocount of 0.1 and projected onto the top 25 principal components using <u>scikit-learn^100^</u>. Unsupervised clustering was performed using <u>Leiden</u> (C++/Python implementation)^21^. Custom Python 3 scripts were used for specific analyses and are available at https://github.com/iosonofabio/lungsc. T Cell receptors were assembled using TraCeR^101^ using the default parameters of the Singularity image. B cell receptors were assembled using BraCeR^102^ with the parameter –IGH_networks, which agreed with our in-house pipeline consisting of Basic^103^ and Change-O^104^. Raw fastq files, count tables, and metadata are available on NCBI’s Gene Expression Omnibus (GEO) website (GSEXXXXX).

### In-situ validation using RNAscope and immunofluorescence (IF)

Embryonic and post-natal mice were euthanized as described above. E18.5 lungs were immediately placed in 10% neutral buffered formalin following dissection. P1, P7, and P21 murine lungs were perfused as described above, and P7 and P21 lungs inflated with 2% low melting agarose (LMT) in 1xPBS, and placed in 10% neutral buffered formalin. Following 20 hours incubation at 4C, fixed lungs were washed twice in 1xPBS and placed in 70% ethanol for paraffin-embedding. In situ validation of genes identified by single cell RNA-seq was performed using the RNAscope Multiplex Fluorescent v2 Assay kit (Advanced Cell Diagnostics) and according to the manufacturer’s protocol. Formalin-fixed paraffin-embedded (FFPE) lung sections (5 μm) were used within a day of sectioning for optimal results. Nuclei were counterstained with DAPI (Life Technology Corp.) and extracellular matrix proteins stained with hydrazide^105^. Opal dyes (Akoya Biosciences) were used for signal amplification as directed by the manufacturer. Images were captured with Zeiss LSM 780 and Zeiss LSM 880 confocal microscopes, using 405nm, 488nm, 560nm and 633nm excitation lasers. For scanning tissue, each image frame was set as 1024×1024 and pinhole 1AiryUnit (AU). For providing Z-stack confocal images, the Z-stack panel was used to set z-boundary and optimal intervals, and images with maximum intensity were processed by merging Z-stacks images. For all both merged signal and split channels were collected.

### Intracellular Flow Cytometry

P1 and P21 male and female murine lungs were isolated as described above. Cells were blocked for 30 minutes with CD16/CD32 (Tonbo Biosciences). For intracellular analyses, cells were stimulated with PMA (50 ng/ml, Sigma-Aldrich), ionomycin (750 ng/mL, Sigma-Aldrich), and GolgiStop (BD Biosciences) for 5 hours, and then surface stained with fluorochrome-conjugated antibodies for 30 minutes: CD3 (clone: 145-2C11, BD Biosciences), CD4 (RM4-5, BD Biosciences), and CD8a (clone: 53-6.7, Biolegend). Cells were then permeabilized with FoxP3 Fixation/Permeabilization Kit (BD Biosciences) as indicated by the manufacturer, and stained for TBET (clone: 4B10, Biolegend), GATA3 (clone: L50-823, BD Biosciences), FOXP3 (clone: FJK-16s, eBioscience), ROR©t (clone: Q31-378, BD Biosciences), IFN© (clone: XMG1.2, Biolegend), IL-4 (clone: 11B11, BD Biosciences), IL-10 (clone: JES5-16E3, Biolegend), and IL-17 (clone: TC11-18H10, Miltenyi Biotec) for 30 minutes. Cells were read using an LSRII flow cytometer using FACSDiva software. Flow data was analyzed using FlowJo (Tree Star Inc.). Flow cytometry analysis for this project was done on instruments in the Stanford Shared FACS Facility.

---

**Supplemental Figure 1.**
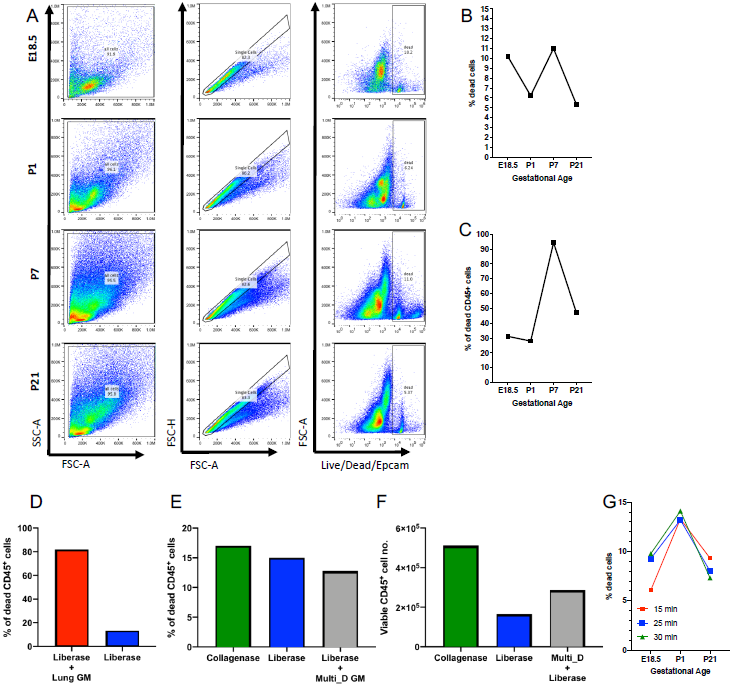
Optimization of lung tissue digestion. (A-C) E18.5, P1, P7 and P21 murine lungs were processed for flow cytometry and the frequency of (B) total and (C) CD45+ (immune) dead cells was assessed. (D-E) Frequency of dead cells of P7 murine lungs was determined by flow cytometry following 30 minute (D) enzymatic digestion with 0.38 mg/mL liberase with manual tituration (Liberase) and/or mechanical disruption using the lung program on the GentleMACS dissociator (Lung GM) or (E) enzymatic digestion with collagenase and manual tituration (Collagenase), Liberase, or 0.38 mg/mL liberase with the Multi_D program on the GentleMACS dissociator (Liberase + Multi_D GM). (F) Number of viable P7 murine lung cells following Collagenase, Liberase, or Liberase + Multi_D GM was quantified. (G) Frequency of murine lung cells at gestational age E18.5, P1, P21 was quantified following incubation with 0.38 mg/mL liberase for 15, 25, and 30 min and manual tituration.

**Supplemental Figure 2.**
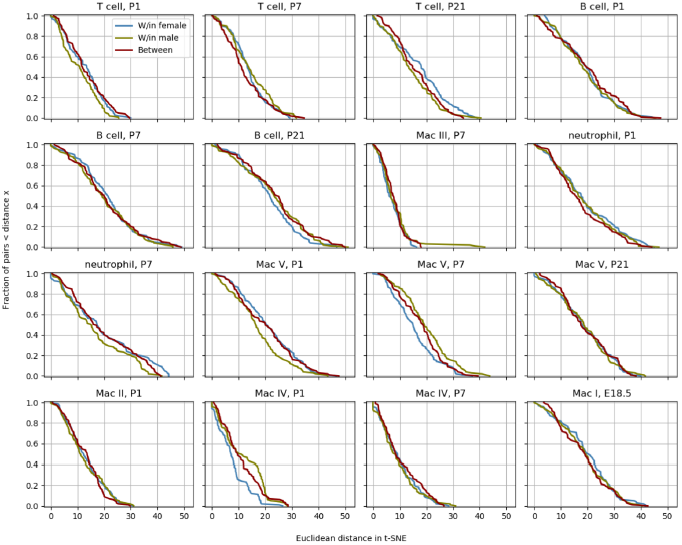
Determination of variation between mice. To quantify whether the different mice contributed spurious variation to the data, a distribution level approach was chosen. For each cell type and time point, 100 pairs of cells from either the same mouse or between different mice were chosen and the distance in tSNE space calculated. The cumulative distributions for those pairs were subsequently plotted to check whether pairs from different animals had a significantly longer distance than cells from the same mouse.

**Supplemental Figure 3.**
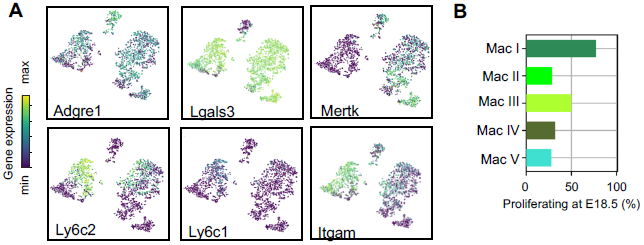
Lineage-defining genes are diffusely expressed across macrophage populations. (A) Graph of percentage of proliferating macrophages in each cluster at E18.5. (B) t-SNE plots of *Adgre1, Lgals3, Mertk, Ly6c2, Ly6c1* and *Itgam* expression across all macrophage clusters between E18.5 and P21.

**Supplemental Figure 4.**
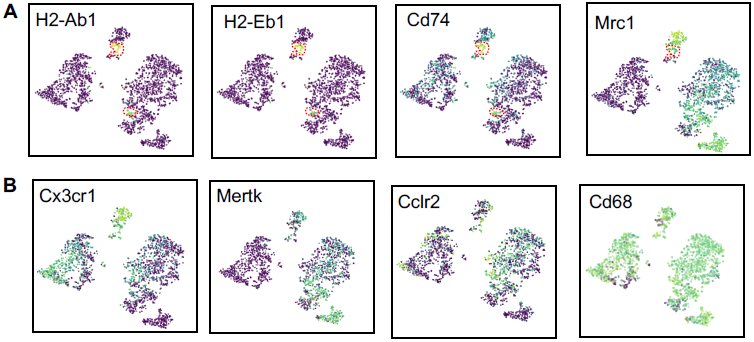
Expression of select genes in Mac IV cluster. (A) t-SNE plots of *H2-Ab1, H2-Eb1, Cd74*, and *Mrcl1* demonstrating differential expression in a portion of Mac IV (dotted red line). (B) t-SNE plots of *Cx3cr1, Mertk, Cclr2, Cd68* demonstrating diffuse expression throughout Mac IV.

**Supplemental Figure 5.**
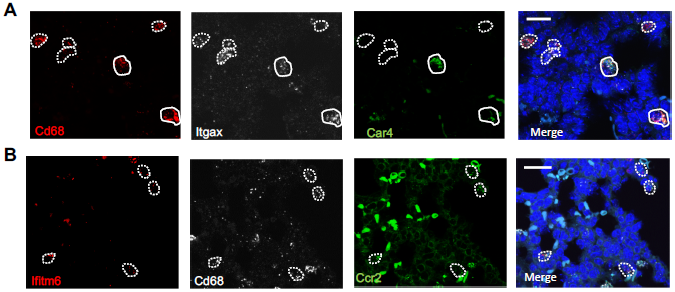
Multiplex In Situ Hybridization to detect specific macrophage clusters. (A)*In situ* hybridization at P1 to detect *Cd68* (red), *Itgax* (white), *Car4* (green) and a merged image. (B) *In situ* hybridization at P1 to detect *Ifitm6* (red), *Cd68* (white) and *Ccr2* (green) and a merged image. Calibration bar=20μm.

**Supplemental Figure 6.**
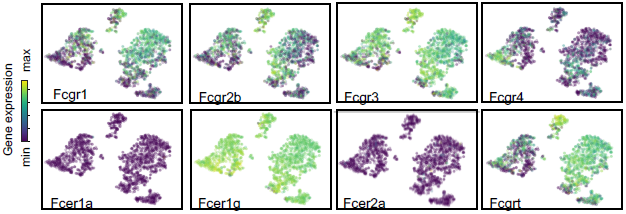
Expression of Fc receptors across macrophage clusters. T-SNE plots of the expression of Fc receptors *Fcgr1, Fcgr2b, Fcgr3, Fcgr4, Fcer1a, Fcer1g, Fcer2a, Fcgrt*.

**Supplemental Figure 7.**
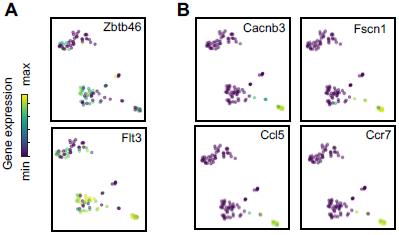
Expression of dendritic cell associated genes. tSNE plots of (A) pan-DC-associated genes *Zbtb46* and *Flt3*, and (B) DCIII-specific associated genes *Cacnb3, Fscn1, Ccl5*, and *Ccr7*.

**Supplemental Figure 8.**
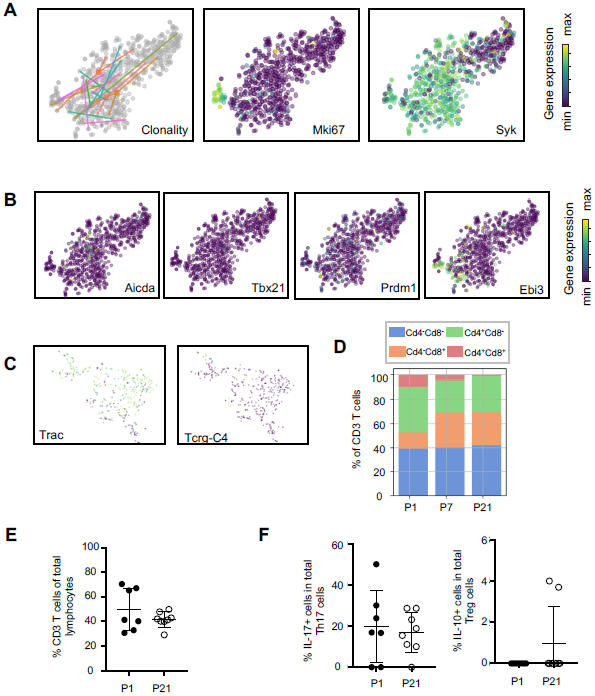
Transcriptional and flow cytometric profiling of lymphocytes. (A) Clonality and t-SNE plots of *Mki67* and *Syk* within the B cell cluster. t-SNE plots of: (B) B cell-associated genes *Aicda, Tbx21, Prdm1*, and *Ebi3*; and (C) T cell-associated genes *Trac* and *Tcrg-C4*. (D) Bar graph of the frequencies of *Cd4+Cd8-, Cd4-Cd8+, Cd4-CD8-*, and *Cd4+Cd8+* T cells at P1, P7, and P21. (E-F) Flow cytometric analyses of murine lung cells isolated at P1 and P21 assessing the frequencies of (E) CD3+ lymphocytes and (F) Il-17-producing Th17 cells and IL-10-producing T regulatory cells.

